# Differential Evasion of Delta and Omicron Immunity and Enhanced Fusogenicity of SARS-CoV-2 Omicron BA.4/5 and BA.2.12.1 Subvariants

**DOI:** 10.1101/2022.05.16.492158

**Authors:** Panke Qu, Julia N. Faraone, John P. Evans, Xue Zou, Yi-Min Zheng, Claire Carlin, Joseph S. Bednash, Gerard Lozanski, Rama K. Mallampalli, Linda J. Saif, Eugene M. Oltz, Peter J. Mohler, Richard J. Gumina, Shan-Lu Liu

## Abstract

The rising case numbers of the SARS-CoV-2 Omicron BA.4, BA.5, and BA.2.12.1 subvariants has generated serious concern about the course of the pandemic. Here we examine the neutralization resistance, infectivity, processing, and fusogenicity of spike from the BA.4/5 and BA.2.12.1 SARS-CoV-2 variants compared with other Omicron subvariants and Delta. Critically, we found that the new Omicron subvariants BA.4/5 and BA.2.12.1 were more resistant to neutralization by mRNA-vaccinated and boosted health care worker sera and Omicron-BA.1-wave patient sera than were the BA.1 and BA.2 variants. Interestingly, Delta-wave patient sera neutralized more efficiently against not only Delta but also BA.4/5 and BA.2.12.1 variants that also contain substitutions at position L452, similar to Delta. The BA.4/5 and BA.2.12.1 variants also exhibited higher fusogenicity, and increased spike processing, dependent on the L452 substitution. These results highlight the key role of the L452R and L452Q mutations in BA.4/5 and BA.2.12.1 subvariants.

## Introduction

The rising frequency of several SARS-CoV-2 Omicron subvariants, especially BA.4/5 and BA.2.12.1, has reignited the new concern about further escape from vaccine- and infection-induced immunity. Since the emergence of SARS-CoV-2 in the human population in 2019, the virus has evolved increased transmissibility (Abdool Karim and de Oliveira, 2021; Scobie et al., 2021; Singh et al., 2021; Washington et al., 2021; Yurkovetskiy et al., 2020) and resistance to vaccine- and infection-induced immunity (Mlcochova et al., 2021; Zhou et al., 2021). The Omicron BA.1 variant was originally discovered in November 2021 and quickly assigned by the World Health Organization as a variant of concern due to its alarming number of mutations (Centers for Disease Control and Prevention, 2022a; Khandia et al., 2022). BA.1 spread rapidly around the globe, causing remarkable levels of breakthrough infection in vaccinated and previously infected individuals because of the continued erosion of immunity (Hoffmann et al., 2022; Pajon et al., 2022; Planas et al., 2022a). Notably, BA.1 demonstrated reduced virulence (Meo et al., 2021) and maintained sensitivity to booster vaccination-induced immunity (Evans et al., 2022a; Pajon et al., 2022).

The recent emergence of several subvariants of Omicron has generated new concerns about vaccine efficacy and the potential for additional waves of the pandemic. BA.1 was responsible for the initial Omicron wave but was supplanted by the more transmissible BA.2 sub-variant (Centers for Disease Control and Prevention, 2022b). BA.2 includes several key mutations that make it distinct from the BA.1 lineage including T19I, L24S, Δ25/27, V213G, T376A, and R408S (outbreak.info, 2022). Critically, BA.2 was able to re-infect patients recently recovered from BA.1 infection (Stegger et al., 2022), and drove additional surges in COVID-19 cases (outbreak.info, 2022). More recently, a derivative of BA.2, namely BA2.12.1, is on the rise in the United States and is characterized by two additional mutations L452Q and S704L (Centers for Disease Control and Prevention, 2022b). Additionally, BA.4 and BA.5 have become the dominant strains in South Africa and are driving a new surge in cases (Network for Genomic Surveillance in South Africa, 2022). The spikes of BA.4 and BA.5 are identical to each other (referred to as BA.4/5 hereafter) but have a unique set of mutations compared to other subvariants (**Fig. S1**). We previously demonstrated significant immune escape by BA.1, BA.1.1, and BA.2 that is overcome by a booster dose of mRNA vaccine (Evans et al., 2022a). In particular, BA.2 could be neutralized by vaccinated sera comparably to BA.1 and was effectively neutralized by sera from those infected during the BA.1 wave (Evans et al., 2022a; Yu et al., 2022). However, the extent of immune escape of the more recently emerged Omicron subvariants, especially BA.4/5 and BA.2.12.1, remains unclear.

To address this question, we examine the neutralizing antibody (nAb) titers in mRNA vaccinated health care workers (HCWs), as well as hospitalized Delta and BA.1-wave patient sera against BA.4/5, and BA.2.12.1 variants, along with BA.1 and BA.2, and compared to the Delta variant and ancestral D614G variant. We also investigate neutralization of the two distinguishing mutations present in BA.2.12.1 that make it distinct from BA.2, i.e., L452Q and S704L. Additionally, we determine and compare the infectivity, processing, and membrane fusion properties of these Omicron sub-lineage variant spike proteins.

## Results

### BA.4/5 and BA.2.12.1 exhibit modestly enhanced resistance to neutralization by 2-dose and 3-dose mRNA-vaccinated HCWs compared to BA.1 and BA.2

We first examined the infectivity of the Omicron subvariants using pseudotyped lentivirus to infect HEK293T-ACE2 or CaLu-3 cells. As we have previously established, the infectivity of BA.1 was slightly higher than D614G (2-fold, p=0.0518) and the Delta variant (5.9-fold, p < 0.001) in HEK293T-ACE2s (**Fig. 1A**), but there was a noted drop in infectivity in CaLu-3 cells (**Fig. 1B**), i.e., 7.5-fold (p<0.0001) and 5.6-fold (p<0.0001) reductions compared to D614G and Delta, respectively (Zeng et al., 2021a). BA.2 showed infectivity comparable to BA.1; however, the remaining subvariants exhibited modestly increased infectivity in 293T-ACE2 cells, with 2.5-fold (p<0.001) and 3.8-fold (p<0.01) increases for BA.4/5 and BA.2.12.1 compared to D614G, respectively (**Fig. 1A**). The L452Q and S704L single mutants showed a 2.9-fold (p<0.001) and 2.3-fold (p<0.01) increase compared to D614G, respectively (**Fig. 1A**). Notably, the infectivity of all Omicron subvariants remained comparably low relative to D614G (p<0.0001) and Delta in CaLu-3 cells (**Fig. 1B**).

**Figure 1:**
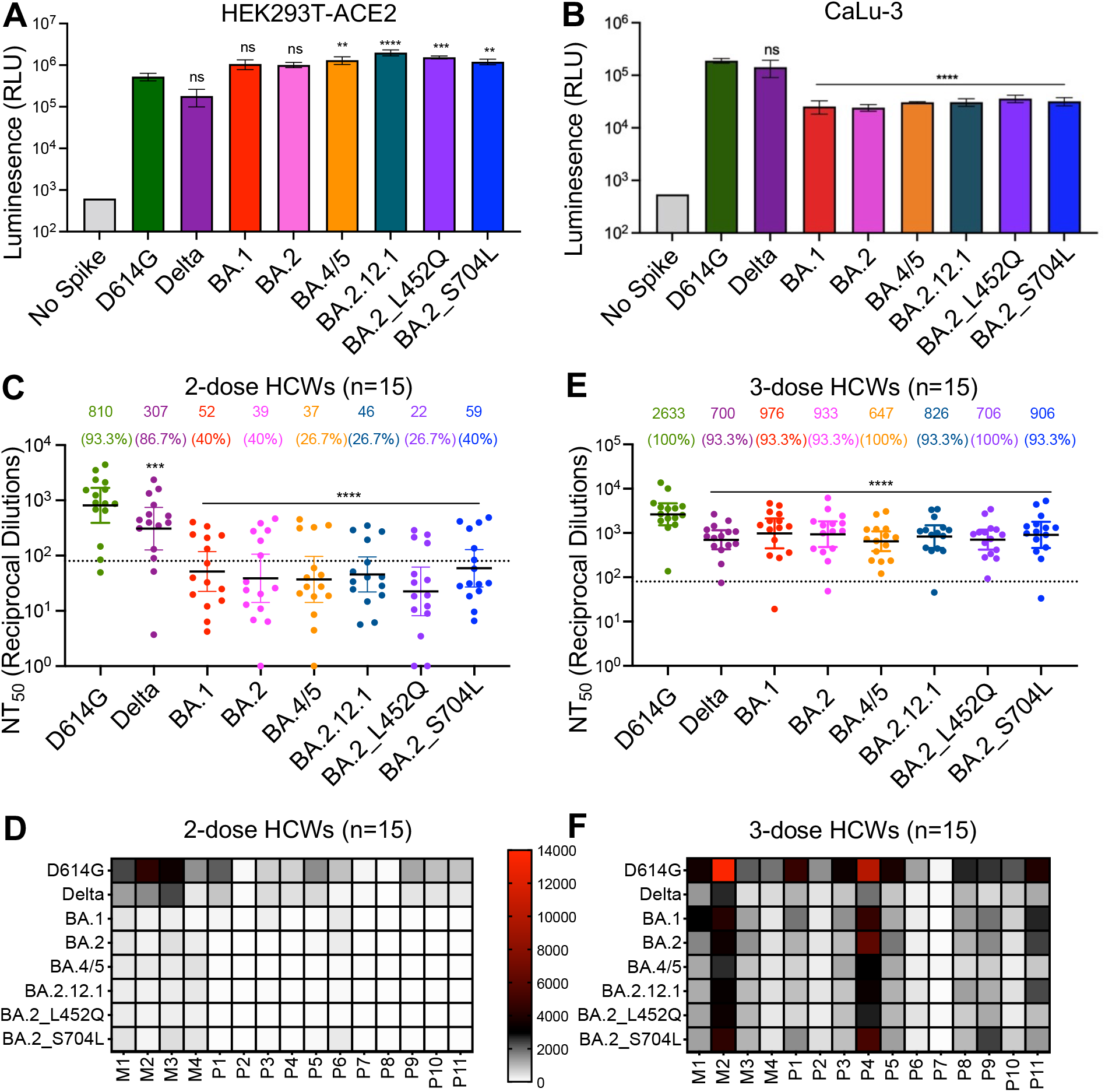
BA.4/5 and BA.2.12.1 subvariants exhibit stronger immune escape than BA.1 and BA.2. (**A**) Infectivity of pseudotyped viruses in HEK293T cells stably expressing ACE2 (HEK293T-ACE2). (**B**) Infectivity of pseudotyped lentivirus in human lung epithelia-derived CaLu-3 cells. Bars in (**A**) and (**B**) represent means ± standard deviation, and significance is determined by one-way repeated measures ANOVA with Bonferroni’s multiple testing correction. Results of at least 3 independent experiments are averaged and shown. (**C**) Sera from 15 HCWs collected 3-4 weeks after second mRNA vaccine dose was used to neutralize pseudotyped virus, and the resulting geometric means of the 50% neutralization titers (NT_50_) are displayed at the top of the graph along with the percent of individuals with NT_50_ values above the limit of detection (NT_50_ = 80; dotted line). (**D**) A heat map showing patient/vaccinee NT_50_ values against each variant for the 2-dose HCW sera. (**E**) Sera from 15 HCWs following homologous mRNA booster vaccination were assessed for nAb titers. Bars in (**C**) and (**E**) represent geometric mean ± 95% confidence interval, and significance relative to D614G is determined by one-way repeated measures ANOVA with Bonferroni’s multiple testing correction. (**F**) A heat map showing patient/vaccinee NT_50_ values against each variant for the 3-dose HCW sera. Patient/vaccinee numbers are identified as “P” for Pfizer/BioNTech BNT162b2 vaccinated/boosted HCW, “M” for Moderna mRNA-1273 vaccinated/boosted HCW. Throughout, p-values are represented as **p < 0.01, ***p < 0.001, ****p < 0.0001, ns, not significant.

We next sought to examine the nAb resistance of the Omicron subvariants in 2-dose mRNA vaccinated HCWs. To address this, we used our previously reported pseudotyped lentivirus neutralization assay (Zeng et al., 2020) to examine the nAb titers in 15 HCWs vaccinated with 2-doses of the Moderna mRNA-1273 (n=4) or Pfizer/BioNTech mRNA-1273 (n=11). We found that the newly emerged BA.4/5 and BA.2.12.1 exhibited similarly potent nAb resistance compared to the BA.1 and BA.2, with nAb titers approximately 20.0-fold (p <0.001) lower than ancestral D614G (**Fig. 1C**). Of note is the greater reduction in nAb titers exhibited by the BA.2_L452Q virus, which was about 36.0-fold lower than that of D614G (p<0.0001) (**Fig. 1C**). Importantly, all these Omicron subvariants exhibited an average of NT_50_ that was below the limit of detection (NT_50_ = 80) (**Fig. 1C and 1D**), consistent with our previous finding that Omicron variants essentially evade neutralization by the 2-dose mRNA vaccination (Evans et al., 2022a).

To determine the sensitivity of the Omicron subvariants BA.4/5 and BA.2.12.1 to sera from recipients of booster mRNA vaccine doses, we examined the nAb titers in 15 HCWs following 3 doses of the Moderna mRNA-1273 (n=4) or Pfizer/BioNTech BNT162b2 (n=11) mRNA vaccines. The nAb titers for all variants dramatically increased after the booster vaccination, as would be expected (**Fig. 1E and 1F**; **Fig. S2A to 2H**). Of note, BA.4/5 and BA.2.12.1 exhibited 4.0-fold (p<0.0001) and 3.2-fold (p<0.0001) reduced NT_50_ compared to D614G, and 31% (p=0.08) and 11% (p=0.90) lower NT_50_ compared to BA.2, respectively (**Fig. 1E**). Again, the resistance phenotype of BA.2_L452Q was stronger than BA.2_S704L, with nAb titers 3.7-fold (p<0.0001) and 2.9-fold (p<0.0001) lower than D614G, respectively (**Fig. 1E**). Importantly, only 1 in 15 of the examined HCWs exhibited an NT_50_ below the limit of detection against all these Omicron subvariants (**Fig. 1F**), demonstrating the effectiveness of booster vaccination in inducing not only markedly higher but also broader nAb titers compared to the 2-dose samples (**Fig. 1D and 1F**). We found that, for both 2-dose and 3-dose mRNA vaccinations, the Modera mRNA-1273 vaccine outperformed the Pfizer/BioNTech BNT162b2 vaccine in the calculated NT_50_ values against all variants, especially for the 2-dose vaccination; however, it should be cautioned that only 4 HCWs in this group received Moderna mRNA-1273 vaccine as compared to 11 HCWs who had received the Pfizer/BioNTech BNT162b2 mRNA vaccine (**Fig. S2I and 2J**).

### Sera from Delta-wave patients potently neutralizes BA.4/5 and BA.2.12.1 subvariants

We next investigated nAb resistance for 18 COVID-19 patients admitted to the intensive care unit (ICU) during the Delta wave of the pandemic. As expected, the nAb titer of Delta patient sera for the Delta variant was among the highest, with even 66.5% higher than D614G (**Fig. 2A**). However, the nAb titers for BA.1 and BA.2 were comparably low, with nAb titers 16.6-fold (p< 0.01) and 17.3-fold (p <0.01) lower than D614G as shown previously (Evans et al., 2022a). Interestingly, we observed an increase in nAb titer of BA.4/5 and BA.2.12.1, with NT_50_ 5.0-fold (p <0.01) and 5.5-fold (p <0.001) lower than D614G, respectively (**Fig. 2A and 2B**). Noticeably, BA.4/5 and BA.2.12.1 exhibited less escape of neutralization by Delta-wave sera, evidenced by fewer NT_50_ values that fell below the limit of detection (**Fig. 2A and 2B**). Interestingly, nAb titers for two single mutants, BA.2_L452Q and BA.2_S704L, decreased relative to BA.4/5 and BA.2.12.1, with BA.2_L452Q 14.9-fold (p <0.0001) and BA.2_S704L 13.8-fold (p <0.001) lower than D614G, respectively (**Fig 2A**). These Delta-wave patients included 12 unvaccinated, 1 vaccinated with 1 dose of the Johnson and Johnson vaccine, 4 patients vaccinated 2 doses of the Pfizer/BioNTech BNT162b2 vaccine, and 1 patient vaccinated with 3 doses of the Moderna mRNA-1273 vaccine. Of note, differences between the NT_50_ values of BA.4/5, BA.2.12.1 and that of two single mutants became less apparent in vaccinated Delta-wave patients compared to unvaccinated Delta-wave patients (**Fig. 2B and 2C**), again indicating that breakthrough infection plus COVID-19 vaccination can offer not only enhanced but also broader protection against the Omicron sub-lineage variants.

**Figure 2:**
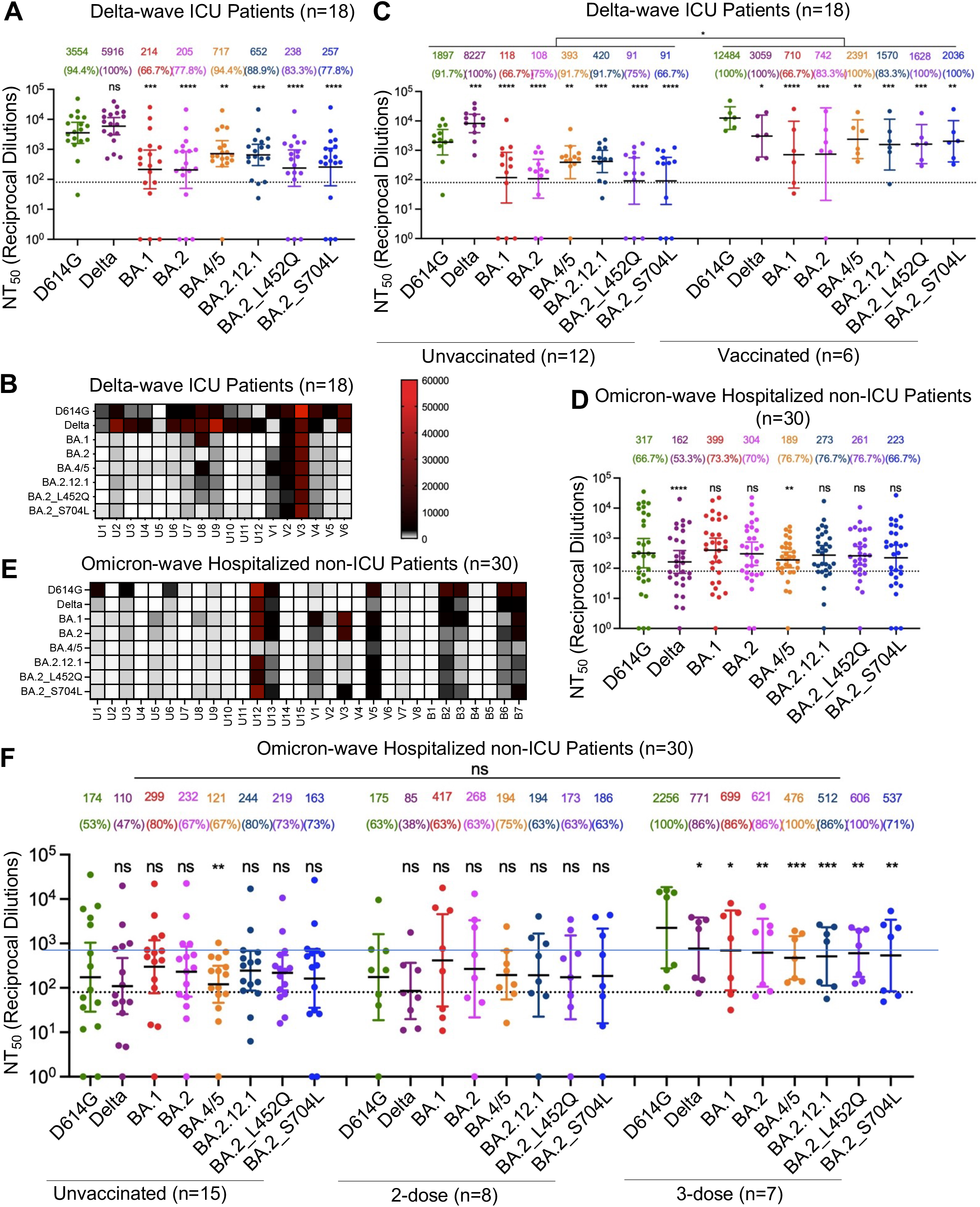
Delta-wave sera more strongly neutralizes BA.4/5 and BA.2.12.1 compared to BA.1 and BA.2. (**A**) Sera from 18 ICU COVID-19 patient samples collected during the Delta wave of the pandemic were assessed for nAb titers. (**B**) A heatmap depicting individual patient NT_50_ values from Delta wave patients. Patients are identified by “U” for unvaccinated and “V” for vaccinated. (**C**) NT_50_ data from (**A**) is plotted by vaccination status with unvaccinated (n=12) and vaccinated (n=6) individuals. (**D**) Sera from 30 hospitalized non-ICU COVID-19 patient samples collected during the BA.1 wave of the pandemic were assessed for nAb titers. (**E**) A heatmap showing patient NT_50_ values against each variant for BA.1-wave patients. Patients are identified by “U” and “V” as described for (**C**) and “B” for 3-dose boosted samples. (**F**) NT_50_ data from (**D**) is plotted by vaccination status with unvaccinated (n=15), 2-dose (n=8), and 3-dose (n=7) individuals depicted. NT_50_ values in (**A, C, D** and **F**) are displayed at the top of plots along with the percentage of patients with NT_50_ values above the limit of detection; bars represent geometric mean ± 95% confidence interval. Significance relative to D614G was determined by one-way repeated measures ANOVA with Bonferroni’s multiple testing correction (**A and D**); significance between vaccination statuses is determined by two-way repeated measures ANOVA with Bonferroni’s multiple testing correction (**C and F**). Throughout, p values are represented as *p < 0.05, **p < 0.01, ***p < 0.001, ****p < 0.0001, ns, not significant.

### BA.4/5 and BA.2.12.1 are remarkably resistant toward neutralization by sera from BA.1-wave hospitalized non-ICU patients

We then sought to characterize nAb resistance for 30 patients hospitalized during the BA.1 wave of Omicron infection. Despite relatively low nAb titers induced by BA.1 infection against all variants tested, the titers of BA.1, BA.2, and D614G were higher than the BA.2-derived sub-lineages (**Fig. 2D**). A modest reduction in neutralization was observed for BA.4/5 and BA.2.12.1, with nAb titers 37.8% (p=0.44) and 10.2% (p>0.999) lower compared to BA.2. Of note, each of the BA.2 single mutants, i.e., BA.2_L452Q and BA.2_S704L, also exhibited an escape comparable to the BA.4/5 and BA.2.12.1 subvariants, with nAb titers 14.1% and 26.6% less than BA.2, respectively (**Fig. 2D**). These BA.1-wave patients included 15 unvaccinated, 4 patients vaccinated with 2 doses of the Pfizer/BioNTech BNT16b2 vaccine, 4 patients vaccinated with 2 doses of the Moderna mRNA-1273 vaccine, and 7 patients vaccinated with 3 doses of the Pfizer/BioNTech BNT16b2 vaccine. We observed that 2 in 30 of BA.1-infected but unvaccinated patients (U12 and U13) developed high nAb titers against all variants, with the notable exception of BA.4/5, yet a 3-dose booster mRNA vaccine significantly improved the nAb titers against all variants examined (**Fig. 2E and 2F**). Overall, these results revealed that the BA.1-wave infection does not seem to offer effective protection against the new emerged sub-lineages BA.4/5 and BA.2.12.1, especially BA.4/5, but a 3-dose booster mRNA vaccination can overcome the deficiency.

### BA.4/5 and BA.2.12.1 subvariants exhibit increased fusogenicity and furin cleavage compared to BA.2

We next determined the propensity for the Omicron subvariant S proteins to mediate membrane fusion. HEK293T-ACE2 cells were co-transfected with the various S protein constructs and GFP and incubated for 24 and 48 hours, and cell-cell fusion was visualized using fluorescence microscopy with size of syncytia formation being quantified (**Fig. 3A and 3B; Fig. S3**). Consistent with our previous data, BA.1 showed markedly lower fusion than the ancestral D614G (4.7-fold) and Delta (11.3-fold) (**Fig. 3B**) (Zeng et al., 2021a). Fusion by BA.2 was observably higher than BA.1 with a 1.5-fold increase (**Fig. 3B**). While still much lower than fusion mediated by Delta (5.2-fold lower for BA.4/5, 5.6-fold lower for BA.2.12.1), the newly emerged Omicron subvariants BA.4/5 and BA.2.12.1 exhibited a higher propensity for fusion, with an average syncytia area 2.1-fold and 1.4-fold higher than BA.1 and BA.2, respectively (**Fig. 3A and 3B**). This phenotype was likely attributed to the L452R or L452Q mutation present in BA.4/5 or BA.1.12.1, as demonstrated by the 2.0-fold increase relative to BA.1 and 1.4-fold increase relative to BA.2 for the BA.2_L452Q single mutant (**Fig. 3B**). Interestingly, BA.2_S704L displayed decreased fusogenicity 1.5-fold lower than BA.2 (**Fig. 3B**).

**Figure 3:**
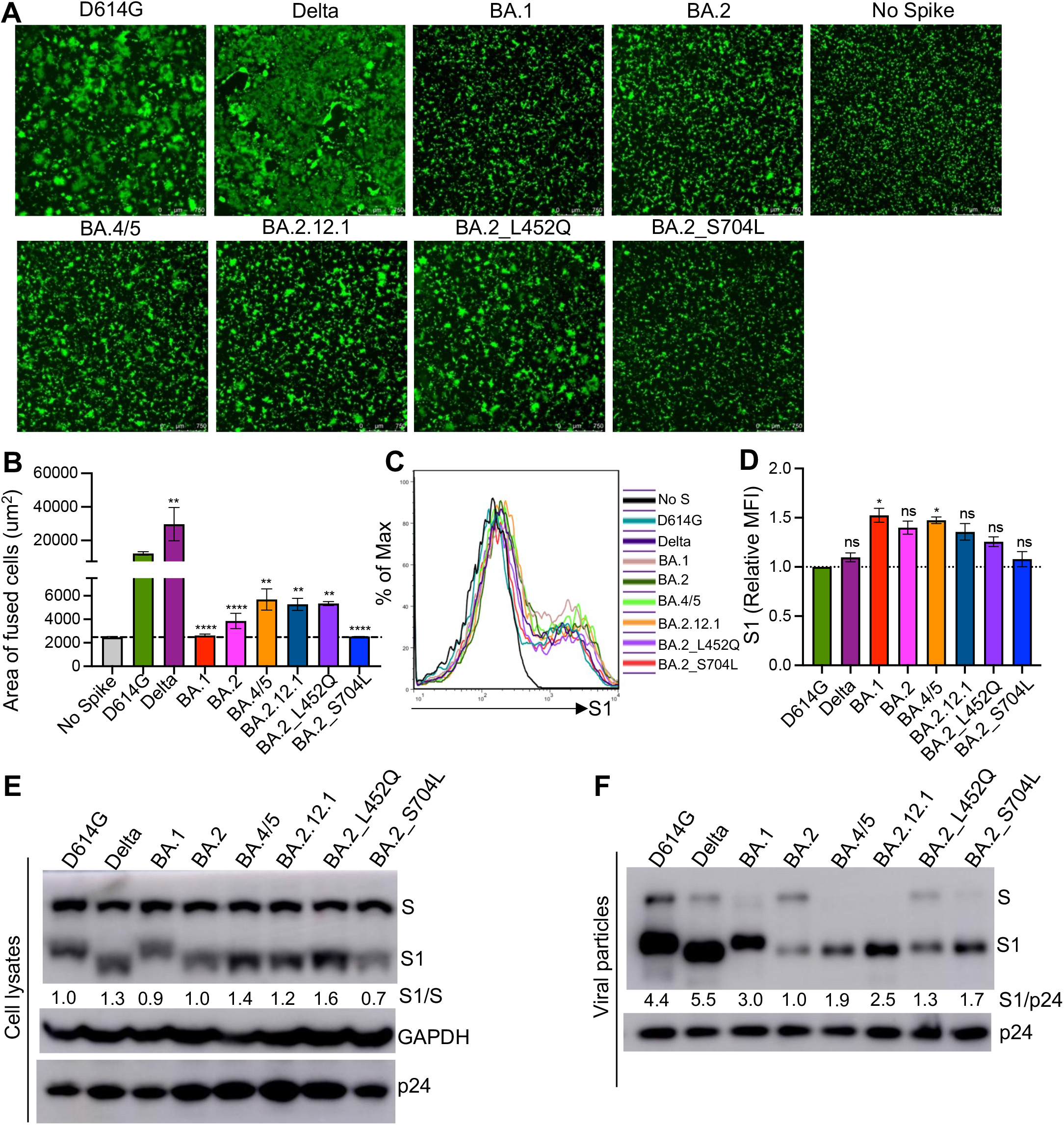
BA.4/5 and BA.2.12.1 S proteins are more fusogenic and more highly processed than BA.2. (**A**) HEK293T cells stably expressing ACE2 were co-transfected with GFP and S constructs and cultured for 24 hours then imaged at 4x magnification to visualize cell-cell fusion. Representative images are depicted. (**B**) Areas of fused cells were quantified using Leica DMi8 confocal microscope analysis software and averaged across 3-4X images, representing 1 biological replicate. (**C**) HEK293T cells were transfected with S constructs and stained for flow cytometry with anti-S1 (T62) to determine S expression. (**D**) Mean fluorescence intensity (MFI) values were normalized to D614G (D614G set to 1.0) and summarized, representing 3 biological replicates. Bars in (**B**) and (**D**) represent means ± standard deviation, and significance was determined by one-way ANOVA with Bonferroni’s multiple testing correction. Throughout, p-values are represented as *p < 0.05, **p < 0.01, ***p < 0.001, ****p < 0.0001, ns, not significant. (**E**) Pseudotyped virus-producing cells were lysed and pseudotyped viruses were purified by ultracentrifugation (**F**) and blots were probed for S1 subunit, HIV-1 capsid (p24), and GAPDH loading control; S cleavage was quantified using NIH ImageJ and by setting the ratio of S1/S of D614G to 1.0 (**E**) and by setting the S1/p24 of D614G to 1.0 for (**F**).

We next examined S expression on the surface of viral producer cells using flow cytometry. Compared to D614G and Delta, BA.1 and BA.2 exhibited slightly higher expression compared to D614G, i.e. 1.5-fold and 1.4-fold relative to D614G, respectively (**Fig. 3C and 3D**). The two new subvariants, i.e., BA.4/5 and BA.2.12.1, exhibited comparable surface expression, with 1.5-fold and 1.4-fold higher surface expression than D614G, respectively (**Fig. 3D**). The BA.2 single mutants, i.e., BA.2_L452Q and BA.2_S704L, both exhibited slightly reduced surface expression compared to BA.4/5 and BA.2.12.1 (**Fig 3C and 3D**).

### Mutation of L452 to R or Q in BA.4/5 or BA.2.12.1 increases spike processing compared to BA.1 and BA.2

We further sought to characterize the processing of the above Omicron subvariant S proteins in cell lysate of virus-producing cells as well as purified viral particles. Lysates from HEK293T cells used to produce the pseudotyped virus were collected and probed for S1 subunit expression versus the full-length S. We have previously demonstrated that BA.1 spike exhibits a lower propensity for furin cleavage, as shown by the reduced ratio of S1/S, in both lysate and purified virions (Zeng et al., 2021a). Here we found that processing of BA.2 S remained comparable to BA.1, but more processing was observed for BA.4/5 and BA.2.12.1, with a 1.4-fold and 1.2-fold higher compared to D614G, respectively (**Fig. 3E**). A similar conclusion could be drawn from the analysis of purified viral particle, wherein BA.4/5, BA.2.12.1 and two single mutants all demonstrated the increased S1 signals in purified virions compared to BA.2 – given similar intensity of p24 in the virions (**Fig. 3F**). In accordance with the fusion results, the increased S processing phenotype in BA.4/5 and BA.2.12.1 was likely attributed to the L452R or L452Q mutation in their receptor binding domain, given that BA.2_L452Q single mutant showed increased S processing in both virus-producing cells and produced virions (**Fig. 3E**). Of note, the processing of BA.2_S704L was slightly reduced in the cells, though not obviously in the virions, ruling out its likely contribution to the BA.2.12.1 phenotype, which was also consistent with its reduced fusogenicity compared to BA.2 (**Fig. 3A, 3B, and 3E**).

## Discussion

In this study, we characterized the infection- and vaccine-induced immunity against the newly emerged Omicron subvariants BA.4/5 and BA.2.12.1 as well as the fusion and furin cleavage properties of their S proteins. We found that two doses of mRNA vaccines, either Pfizer or Moderna, were not sufficient for neutralization of any of the Omicron subvariants, which agrees with trends reported by ourselves and others on BA.1 (Evans et al., 2022a; Pérez-Then et al., 2022; Planas et al., 2022b; Schmidt et al., 2022; Wang et al., 2022; Xia et al., 2022) and BA.2 (Evans et al., 2022a; Yu et al., 2022). Critically, booster vaccination can provide sufficient nAb titers against the subvariants but with approximately 35% drop in nAb titer against BA.4/5 compared to BA.1 and BA.2. Our data suggest that mutations at residue 452 (L452R in BA.4/5 and L452Q in BA.2.12.1) may be one of the key drivers of this neutralization resistance phenotype. L452R is also a defining mutation of the Delta variant that has been previously established as granting neutralization resistance (Patrick et al., 2022; Zhang et al., 2022b). Not surprisingly, the new Omicron subvariants containing mutations at 452 are more effectively neutralized by Delta-wave ICU patients. Alarmingly, however, these two new Omicron subvariants have started to evade the humoral immunity induced by Omicron itself, evidenced by the poor efficacy of BA.1-wave sera in neutralizing the new subvariants. While further investigations are warranted with regards to the molecular mechanisms, these results further emphasize the importance of booster vaccination, as opposed to natural BA.1-wave infection-induced immunity, in neutralizing these new subvariants.

We demonstrated a notable increase in infectivity for these new subvariants in HEK293T (human kidney) cells stably expressing ACE2 yet dramatic decreases in infectivity in human lung epithelial CaLu-3 cells. These results are consistent with our previous report on BA.1 containing R346K mutation (Zeng et al., 2021a), together suggesting that the new Omicron variants may not replicate with high efficiency in the lower airway epithelia derived cells compared to other variants of concern. Low viral load in the lungs compared to the nasal airway has since been demonstrated for BA.1 (Brüssow, 2022; Hui et al., 2022), and poor replication in CaLu-3 cells rather than in primary human nasal epithelial cells has been reported for BA.2 (Yamasoba et al., 2022). Our data could suggest that the BA.4/5 and BA.2.12.1 variants may retain the reduced pathogenicity of the BA.1 variant. However, we also find that BA.4/5, and BA.2.12.1., along with BA.2, exhibit apparently enhanced fusogenicity compared to BA.1. We proposed previously that the attenuation of fusion in BA.1 containing R346K mutation was likely a fitness cost to more effectively evade neutralization (Zeng et al., 2021a). However, given that the extent of syncytia formation has been correlated with pathogenicity in past studies (Braga et al., 2021; Zhang et al., 2021), our results may suggest potentially enhanced transmission and pathogenesis of the BA.4/5 and BA.2.12.1 Omicron subvariants, which appears to be supported by some recent reports (Yamasoba et al., 2022; Zhang et al., 2022a). Critically, we find that, as with neutralization, the L452Q substitution plays a key role in fusion, further emphasizing the importance of this mutation. The increased fusogenicity could be associated with enhanced S cleavage, as observed for BA.4/5, BA.2.12.1, and BA.2_L452Q compared to BA.2. However, more studies are needed to determine the molecular mechanism for the discrepancy between fusion and S process of BA.2 and BA.1

Overall, our findings emphasize the importance of booster vaccination in control of the continually evolving Omicron variants. The waning extent of neutralization by vaccinated sera suggests (Evans et al., 2022a, 2022b) that development of more broadly neutralizing vaccines may be required in the near future to maintain current levels of control. The differences exhibited in neutralization by BA.1-wave vs. Delta-wave sera demonstrate a strong selective pressure being imposed by immunity induced by previous Omicron subvariants, evidenced by the re-appearance of L452 substitutions in BA.4/5 and BA.2.12.1. The continued evolution of the virus in this manner could also contribute to the waning nature of vaccinated immunity. Critically, these two new subvariants of Omicron exhibit increased fusogenicity, which may suggest enhanced transmission, cell-to-cell spread, and pathogenesis (Zeng et al., 2021a, 2022). The rapid manner in which SARS-CoV-2 continues to evolve, including the recombination between variants of concern resulting in Deltacron and other new variants (Colson et al., 2022; Evans et al., 2022c; Lacek et al.; Ou et al., 2022) demonstrates the importance of monitoring emerging variants and understanding the implications of S protein biology on transmissibility, pathogenicity, and immunity.

## Acknowledgements

We thank the NIH AIDS Reagent Program and BEI Resources for providing important reagents for this work. We also thank the Clinical Research Center/Center for Clinical Research Management of The Ohio State University Wexner Medical Center and The Ohio State University College of Medicine in Columbus, Ohio, specifically Francesca Madiai, Dina McGowan, Breona Edwards, Evan Long, and Trina Wemlinger, for logistics, collection and processing of samples. In addition, we thank Sarah Karow, Madison So, Preston So, Daniela Farkas, and Finny Johns in the clinical trials team of The Ohio State University for sample collection and other supports.

## Author Contributions

S.-L.L. conceived and directed the project. P.Q. performed most of the experiments. J.N.F, X.Z., J.P.E. assisted in experiments and contributed data processing and analyses. C.C., J.S.B., G.L., R.K.M., R.J.G. provided clinical samples. P.Q., J.N.F., J.P.E., and S.-L.L. wrote the paper. P.J.M. facilitated shipping of the original Omicron construct. Y.-M.Z, L.J.S., E.M.O., P.J.M., and R.J.G. provided insightful discussion and revision of the manuscript.

## Star Methods

### RESOURCE AVAILABILITY

#### Lead contact

Further information and requests for resources and reagents should be directed to the lead contact, Dr. Shan-Lu Liu.

#### Materials availability

Plasmids generated in this study are available upon request made to the lead contact.

#### Data and code availability

- NT_50_ values and de-identified patient information will be deposited to the National Cancer Institute SeroNet Coordinating Center. Additionally, NT_50_ values and de-identified patient information reported in this paper will be shared by the lead contact upon request.
- This paper does not report original code.
- Any additional information required to reanalyze the data reported in this paper is available from the lead contact upon request

### EXPERIMENTAL MODEL AND SUBJECT DETAILS

#### Patient Information

Two-dose vaccinated HCW samples were collected under approved IRB protocols (2020H0228 and 2020H0527). Demographic information was self-reported, and all subjects provided informed consent. Sera were collected 3-4 weeks post-second vaccine dose for 15 HCWs (7 female and 8 male; median age 37; age range 31-56), which included 4 Moderna mRNA-1273 and 11 Pfizer/BioNTech BNT162b2 vaccinated HCWs.

Boosted HCW samples were additionally collected under the same IRB protocols. Demographic information was self-reported, and all subjects provided informed consent. Sera were collected 1-11 weeks post-homologous booster vaccine dose for 15 HCWs (7 female and 8 male; median age 37; age range 22-48), which included 4 Moderna mRNA-1273 and 11 Pfizer/BioNTech BNT162b2 vaccinated HCWs. Note that 12 HCWs had both post-second dose samples and post-booster dose samples analyzed.

Delta-wave ICU patient samples were collected under an approved IRB protocol (2020H0175). Demographic information was self-reported, and all subjects provided informed consent. Plasma samples were collected 3 days after ICU admission for 18 Delta-wave patients (6 female and 12 male; median age 60; age range 22-87; 4 African American/Black non-Hispanic or Latino, 1 White Hispanic or Latino, and 13 White non-Hispanic or Latino). Where detectable, the variant of SARS-CoV-2 infecting the ICU patients was confirmed by viral RNA extraction on nasal swabs with QIAamp MinElute Virus Spin kit followed by RT-PCR (CDC N1 F: 5’-GACCCCAAAATCAGCGAAAT-3’; CDC N1 R: 5’-TCTGGTTACTGCCAGTTGAATCTG-3’; CDC N2 F: 5’-TTACAAACATTGGCCGCAAA-3’; CDC N2 R: 5’-GCGCGACATTCCGAAGAA-3’) and Sanger sequencing to identify the variant. In total, 5/18 patients were confirmed to have been infected with Delta. Additionally, these Delta-wave patients included 1 patient vaccinated with 1 dose of the Johnson & Johnson vaccine, 4 patients vaccinated with 2 doses of the Pfizer/BioNTech BNT162b2 vaccine, and 1 patient vaccinated with 3 doses of the Moderna mRNA-1273 vaccine.

Omicron-wave hospitalized patient samples were collected under an approved IRB (2020H0527). Demographic information was self-reported, and all subjects provided informed consent. Sera were collected 1-8 days after hospitalization for 30 COVID-19 patients (11 female and 19 male; median age 62; age range 28-78) admitted in late January and February of 2022. These included 15 unvaccinated patients. Additionally, 8 patients were vaccinated with two doses of the Pfizer/BioNTech BNT16b2 vaccine (n = 4) or Moderna mRNA-1273 vaccine (n = 4), and sample collection occurred 5-11 months (median 9 months) after 2nd vaccine dose. Finally, 7 patients were vaccinated with three doses of the Pfizer/BioNTech BNT162b2 vaccine and sample collection occurred 2-6 months (median 5 months) after booster vaccine administration.

#### Cell lines and maintenance

All cell lines were maintainted in 5% CO_2_ and at 37°C. HEK293T (ATCC CRL-11268, RRID: CVCL_1926) and HEK293T-ACE2 (BEI NR-52511, RRID: CVCL_A7UK) cells were maintained in DMEM (Gibco, 11965-092) supplemented with 10% FBS (Sigma, F1051) and 1% penicillin-streptomycin (HyClone, SV30010). CaLu-3 cells (RRID: CVCL_0609) were maintained in EMEM (ATCC 30-2003) supplemented with 10% FBS and 1% penicillin-streptomycin.

### METHOD DETAILS

#### Plasmids

Pseudotyped virus was produced using a pNL4-3-inGluc lentivirus vector which contains a ΔENV HIV-1 backbone bearing a *Gaussia* luciferase reporter gene (Goerke et al., 2008; Zeng et al., 2020). GenScript Biotech (Piscataway, NJ) produced and cloned SARS-CoV-2 spike constructs with N- and C-terminal flag tags using Kpn I and BamH I restriction enzyme cloning into a pcDNA3.1 vector.

#### Pseudotyped lentivirus production and infectivity

Lentiviral pseudotypes were produced as previously described (Evans et al., 2021). pNL4-3-inGluc and spike constructs were transfected into HEK-293T cells in a 2:1 ration using polyethylenimine transfection. Pseudotyped virus was harvested 24, 48, and 72 hrs after transfection. Pseudotyped virus for each SARS-CoV-2 spike, produced in parallel, were used to infect target HEK293T-ACE2 or CaLu-3 cells. *Gaussia* luciferase activity was assessed 48 hrs after infection by combining *Gaussia* luciferase substrate (0.1M Tris pH 7.4, 0.3M sodium ascorbate, 10 μM coelenterazine) with cell culture media. Luminescence was immediately measured by a BioTek Cytation5 plate reader.

#### Virus neutralization assay

Pseudotyped lentivirus neutralization assays were performed as previously described (Evans et al., 2021; Zeng et al., 2020, 2021b, 2021c). HCW serum or ICU patient plasma was serially diluted 4-fold. Equal amounts of pseudotyped lentivirus bearing SARS-CoV-2 S protein was added to the diluted serum. Final dilutions of 1:80, 1:320, 1:1280, 1:5120, 1:20480, and no serum control for HCW and BA.1-wave samples. Delta-wave samples were treated with Triton X-100 to inactivate virus. To prevent Triton X-100 toxicity from impacting the assay, final dilutions for Delta-wave serum were 1:1280, 1:2560, 1:5120, 1:10240, and 1:20480. Virus and serum were incubated for 1 hr at 37°C and then added to HEK293T-ACE2 cells for infection by neutralized virus. Gaussia luciferase output by infected cells was measured 48 and 72 hrs after infection by taking 20 μL of infected cell culture media and adding 20 μL of Gaussia luciferase substrate. Luminescence was measured immediately using a BioTek Cytation5 plate reader. NT_50_ values were determined by least-squares-fit, non-linear regression in GraphPad Prism 9 (San Diego, CA). Heat maps with NT_50_ generated by GraphPad Prism 9.

#### Spike detection by flow cytometry

HEK293T cells used to produce pseudotyped vectors were harvested and fixed 72 hrs after transfection. Cells were dissociated in PBS + 5mM EDTA at 37°C for 30 min. Once dissociated, cells were fixed in 4% formaldehyde diluted in 1X PBS and stained with primary antibody anti-S1 (Sino Biological, 40150-T62). Cells were then stained with secondary antibody anti-rabbit-IgG-FITC (Sigma, F9887) and processed by a Life Technologies Attune NxT flow cytometer. Results were analyzed using FlowJo v7.6.5 (Ashland, OR).

#### Syncytia Formation Assay

HEK293T-ACE2 cells were co-transfected with variant spikes of interest and a GFP expression plasmid. Cells were incubated for 24 and 48 hours and images were taken at 4x magnification on a Leica DMi8 confocal microscope. Three images were taken for each sample and used to quantify total area of fused cells (syncytia) using the Leica microscope analysis software. A representative image was selected for each.

#### Spike incorporation into pseudotyped virus

Purification of pseudotyped particles was performed using ultracentrifugation through a 20% sucrose cushion. The resulting pellet was resuspended in SDS-PAGE loading buffer in preparation for western blotting. Additionally, the cell lysate of virus producing cells was collected 72 hours post transfection through lysis with RIPA buffer (50 mM Tris pH 7.5, 150 mM NaCl, 1 mM EDTA, Nonidet P-40, 0.1% SDS) supplemented with protease inhibitor (Sigma, P8340) on ice for 30 min Purification and lysate samples were run on a 10% acrylamide SDS-PAGE gel and transferred to a PVDF membrane. Membranes were blotted with anti-S1 (Sino Biological, 40150-T62), anti-p24 (Abcam, ab63917; NIH ARP-1513), and anti-GAPDH (Santa Cruz Biotech, sc-47724). Anti-mouse-IgG-Peroxidase (Sigma, A5278) and anti-rabbit-IgG-HRP (Sigma, A9169) were used as secondary antibodies accordingly. Blots were imaged with Immobilon Crescendo Western HRP substrate (Millipore, WBLUR0500) on a GE Amersham Imager 600.

### QUANTIFICATION AND STATISTICAL ANALYSIS

All statistical analysis was performed using GraphPad Prism 9 and are described in the figure legends. NT_50_ values were determined by least-squares fit non-linear regression in GraphPad Prism 9. Throughout, n refers to subject number and bars represent either ± standard deviation (**Fig 1A and B, Fig 2B and D**) or geometric means ± 95% confidence intervals (**Fig 1C and E, Fig2A, C, D, F and Fig S2I and J**).Generally, comparisons between multiple groups were made using a one-way repeated measures ANOVA with Bonferroni post-test (**Fig 1A, B, C and E, 2A and D, 3B and D**). Comparisons between treatment groups were made using a two-way repeated measures ANOVA with Bonferroni post-test (**Fig 2C and F, Fig S2I and J**). Comparisons between two-groups were made using a paired two-tailed student’s t-test with Welch’s correction (**Fig S2A-H**). Analysis of the influence of sex on neutralization capacity could not be performed due to small sample size—confounding variables including vaccination status, vaccine type, and time since vaccination would skew such an analysis. Infectivity was quantified using luminescence readings measured by a BioTek Cytation5 plate reader. Three readings were taken and averaged for each of the three replicates. Quantification of syncytia was performed using the Leica DMi8 confocal microscope analysis software to identify the edges of syncytia and quantify area. Surface staining was quantified with a Life Technologies Attune NxT flow cytometer. Results were processed using FlowJo v7.6.5. Western Blot quantification was performed using NIH ImageJ and by setting the ratio of S1/S and S1/p24 of BA.2 to 1.00.

## References

Abdool Karim, S.S., and de Oliveira, T. (2021). New SARS-CoV-2 Variants — Clinical, Public Health, and Vaccine Implications. New England Journal of Medicine 384, 1866–1868. https://doi.org/10.1056/nejmc2100362.

Braga, L., Ali, H., Secco, I., Chiavacci, E., Neves, G., Goldhill, D., Penn, R., Jimenez-Guardeño, J.M., Ortega-Prieto, A.M., Bussani, R., et al. (2021). Drugs that inhibit TMEM16 proteins block SARS-CoV-2 spike-induced syncytia. Nature 594, 88–93. https://doi.org/10.1038/s41586-021-03491-6.

Brüssow, H. (2022). COVID-19: Omicron – the latest, the least virulent, but probably not the last variant of concern of SARS-CoV-2. Microbial Biotechnology https://doi.org/10.1111/1751-7915.14064.

Centers for Disease Control and Prevention (2022a). SARS-CoV-2 Variant Classifications and Definitions.

Centers for Disease Control and Prevention (2022b). Variant Proportions.

Colson, P., Fournier, P., Delerce, J., Million, M., Bedotto, M., Houhamdi, L., Yahi, N., Bayette, J., Levasseur, A., Fantini, J., et al. (2022). Culture and identification of a “Deltamicron” SARS-CoV-2 in a three cases cluster in southern France. Journal of Medical Virology https://doi.org/10.1002/jmv.27789.

Evans, J.P., Zeng, C., Carlin, C., Lozanski, G., Saif, L.J., Oltz, E.M., Gumina, R.J., and Liu, S.-L. (2021). Loss of Neutralizing Antibody Response to mRNA Vaccination against SARS-CoV-2 2 Variants: Differing Kinetics and Strong Boosting by Breakthrough Infection 3 4 5. BioRxiv https://doi.org/10.1101/2021.12.06.471455.

Evans, J.P., Zeng, C., Qu, P., Faraone, J., Zheng, Y.-M., Carlin, C., Bednash, J.S., Zhou, T., Lozanski, G., Mallampalli, R., et al. (2022a). Neutralization of SARS-CoV-2 Omicron Sub-lineages BA.1, BA.1.1, and BA.2. Cell Host & Microbe https://doi.org/10.1016/j.chom.2022.04.014.

Evans, J.P., Zeng, C., Carlin, C., Lozanski, G., Saif, L.J., Oltz, E.M., Gumina, R.J., and Liu, S.-L. (2022b). Neutralizing antibody responses elicited by SARS-CoV-2 mRNA vaccination wane over time and are boosted by breakthrough infection.

Evans, J.P., Qu, P., Zeng, C., Zheng, Y.-M., Carlin, C., Bednash, J.P., Lozanski, G., Mallampalli, R., Saif, L.J., Oltz, E.M., et al. (2022c). Neutralization of the SARS-CoV-2 Deltacron and BA.3 Variants. New England Journal of Medicine. In Press.

Goerke, A.R., Loening, A.M., Gambhir, S.S., and Swartz, J.R. (2008). Cell-free metabolic engineering promotes high-level production of bioactive Gaussia princeps luciferase. Metabolic Engineering 10, 187–200. https://doi.org/10.1016/j.ymben.2008.04.001.

Hoffmann, M., Krüger, N., Schulz, S., Cossmann, A., Rocha, C., Kempf, A., Nehlmeier, I., Graichen, L., Moldenhauer, A.S., Winkler, M.S., et al. (2022). The Omicron variant is highly resistant against antibody-mediated neutralization: Implications for control of the COVID-19 pandemic. Cell 185, 447–456.e11. https://doi.org/10.1016/j.cell.2021.12.032.

Hui, K.P.Y., Ho, J.C.W., Cheung, M. chun, Ng, K. chun, Ching, R.H.H., Lai, K. ling, Kam, T.T., Gu, H., Sit, K.Y., Hsin, M.K.Y., et al. (2022). SARS-CoV-2 Omicron variant replication in human bronchus and lung ex vivo. Nature 603, 715–720. https://doi.org/10.1038/s41586-022-04479-6.

Khandia, R., Singhal, S., Alqahtani, T., Kamal, M.A., El-Shall, N.A., Nainu, F., Desingu, P.A., and Dhama, K. (2022). Emergence of SARS-CoV-2 Omicron (B.1.1.529) variant, salient features, high global health concerns and strategies to counter it amid ongoing COVID-19 pandemic. Environmental Research 209. https://doi.org/10.1016/j.envres.2022.112816.

Lacek, K.A., Rambo-Martin, B.L., Batra, D., Zheng, X.-Y., Sakaguchi, H., Peacock, T., Keller, M., Wilson, M.M., Sheth, M., Davis, M.L., et al. Identification of a Novel SARS-CoV-2 Delta-Omicron Recombinant Virus in the United States. https://doi.org/10.1101/2022.03.19.484981.

Meo, S.A., Meo, A.S., Al-Jassir, F.F., and Klonoff, D.C. (2021). Omicron SARS-CoV-2 new variant: global prevalence and biological and clinical characteristics. Eur Rev Med Pharmacol Sci 8012–8018.

Mlcochova, P., Kemp, S., Dhar, M.S., Papa, G., Meng, B., Ferreira, I.A.T.M., Datir, R., Collier, D.A., Albecka, A., Singh, S., et al. (2021). SARS-CoV-2 B.1.617.2 Delta variant replication and immune evasion. Nature https://doi.org/10.1038/s41586-021-03944-y.

Network for Genomic Surveillance in South Africa (2022). SARS-CoV-2 Sequencing Update 29 April 2022 (Wiley).

Ou, J., Lan, W., Wu, X., Zhao, T., Duan, B., Yang, P., Ren, Y., Quan, L., Zhao, W., Seto, D., et al. (2022). Tracking SARS-CoV-2 Omicron diverse spike gene mutations identifies multiple inter-variant recombination events. Signal Transduction and Targeted Therapy 7, 138. https://doi.org/10.1038/s41392-022-00992-2.

Outbreak.info (2022). Omicron Variant Report.

Pajon, R., Doria-Rose, N.A., Shen, X., Schmidt, S.D., O’Dell, S., McDanal, C., Feng, W., Tong, J., Eaton, A., Maglinao, M., et al. (2022). SARS-CoV-2 Omicron Variant Neutralization after mRNA-1273 Booster Vaccination. New England Journal of Medicine 386, 1088–1091. https://doi.org/10.1056/nejmc2119912.

Patrick, C., Upadhyay, V., Lucas, A., and Mallela, K.M.G. (2022). Biophysical fitness landscape of the SARS-CoV-2 Delta variant receptor binding domain. Journal of Molecular Biology 167622. https://doi.org/10.1016/j.jmb.2022.167622.

Pérez-Then, E., Lucas, C., Monteiro, V.S., Miric, M., Brache, V., Cochon, L., Vogels, C.B.F., Malik, A.A., de la Cruz, E., Jorge, A., et al. (2022). Neutralizing antibodies against the SARS-CoV-2 Delta and Omicron variants following heterologous CoronaVac plus BNT162b2 booster vaccination. Nature Medicine 28, 481–485. https://doi.org/10.1038/s41591-022-01705-6.

Planas, D., Saunders, N., Maes, P., Guivel-Benhassine, F., Planchais, C., Buchrieser, J., Bolland, W.H., Porrot, F., Staropoli, I., Lemoine, F., et al. (2022a). Considerable escape of SARS-CoV-2 Omicron to antibody neutralization. Nature 602, 671–675. https://doi.org/10.1038/s41586-021-04389-z.

Planas, D., Saunders, N., Maes, P., Guivel-Benhassine, F., Planchais, C., Buchrieser, J., Bolland, W.H., Porrot, F., Staropoli, I., Lemoine, F., et al. (2022b). Considerable escape of SARS-CoV-2 Omicron to antibody neutralization. Nature 602, 671–675. https://doi.org/10.1038/s41586-021-04389-z.

Schmidt, F., Muecksch, F., Weisblum, Y., da Silva, J., Bednarski, E., Cho, A., Wang, Z., Gaebler, C., Caskey, M., Nussenzweig, M.C., et al. (2022). Plasma Neutralization of the SARS-CoV-2 Omicron Variant. New England Journal of Medicine 386, 599–601. https://doi.org/10.1056/nejmc2119641.

Scobie, H.M., Johnson, A.G., Suthar, A.B., Severson, R., Alden, N.B., Balter, S., Daniel Bertolino,;, Blythe, D., Brady, S., Cadwell, B., et al. (2021). Monitoring Incidence of COVID-19 Cases, Hospitalizations, and Deaths, by Vaccination Status — 13 U.S. Jurisdictions, April 4–July 17, 2021. Morbidity and Mortality Weekly Report https://doi.org/10.1101/2021.08.11.21261885v1.

Singh, J., Rahman, S.A., Ehtesham, N.Z., Hira, S., and Hasnain, S.E. (2021). SARS-CoV-2 variants of concern are emerging in India. Nature Medicine 27, 1131–1133. https://doi.org/10.1038/s41591-021-01397-4.

Stegger, M., Marie Edslev, S., Niklaus Sieber, R., Cäcilia Ingham, A., Ng, L., Eric Tang, M.-H., Alexandersen, S., Fonager, J., Legarth, R., Utko, M., et al. (2022). Occurrence and significance of Omicron BA.1 infection followed by 1 BA.2 reinfection 2 3. MedRxiv https://doi.org/10.1101/2022.02.19.22271112.

Wang, K., Jia, Z., Bao, L., Wang, L., Cao, L., Chi, H., Hu, Y., Li, Q., Zhou, Y., Jiang, Y., et al. (2022). Memory B cell repertoire from triple vaccinees against diverse SARS-CoV-2 variants. Nature 603, 919–925. https://doi.org/10.1038/s41586-022-04466-x.

Washington, N.L., Gangavarapu, K., Zeller, M., Bolze, A., Cirulli, E.T., Schiabor Barrett, K.M., Larsen, B.B., Anderson, C., White, S., Cassens, T., et al. (2021). Emergence and rapid transmission of SARS-CoV-2 B.1.1.7 in the United States. Cell 184, 2587–2594.e7. https://doi.org/10.1016/j.cell.2021.03.052.

Xia, H., Zou, J., Kurhade, C., Cai, H., Yang, Q., Cutler, M., Cooper, D., Muik, A., Jansen, K.U., Xie, X., et al. (2022). Neutralization and durability of 2 or 3 doses of the BNT162b2 vaccine against Omicron SARS-CoV-2. Cell Host and Microbe 30, 485–488.e3. https://doi.org/10.1016/j.chom.2022.02.015.

Yamasoba, D., Kimura, I., Nasser, H., Morioka, Y., Nao, N., Ito, J., Uriu, K., Tsuda, M., Zahradnik, J., Shirakawa, K., et al. (2022). Virological characteristics of SARS-CoV-2 BA.2 variant. BioRxiv https://doi.org/10.1101/2022.02.14.480335.

Yu, J., Collier, A.-R.Y., Rowe, M., Mardas, F., Ventura, J.D., Wan, H., Miller, J., Powers, O., Chung, B., Siamatu, M., et al. (2022). Comparable Neutralization of the SARS-CoV-2 Omicron BA.1 and BA.2 Variants Authors for Print Edition. MedRxiv https://doi.org/10.1101/2022.02.06.22270533.

Yurkovetskiy, L., Wang, X., Pascal, K.E., Tomkins-Tinch, C., Nyalile, T.P., Wang, Y., Baum, A., Diehl, W.E., Dauphin, A., Carbone, C., et al. (2020). Structural and Functional Analysis of the D614G SARS-CoV-2 Spike Protein Variant. Cell 183, 739–751.e8. https://doi.org/10.1016/j.cell.2020.09.032.

Zeng, C., Evans, J.P., Pearson, R., Qu, P., Zheng, Y.M., Robinson, R.T., Hall-Stoodley, L., Yount, J., Pannu, S., Mallampalli, R.K., et al. (2020). Neutralizing antibody against SARS-CoV-2 spike in COVID-19 patients, health care workers, and convalescent plasma donors. JCI Insight 5. https://doi.org/10.1172/jci.insight.143213.

Zeng, C., Evans, J.P., Qu, P., Faraone, J., Zheng, Y.-M., Carlin, C., Bednash, J.S., Zhou, T., Lozanski, G., Mallampalli, R., et al. (2021a). Neutralization and Stability of SARS-CoV-2 Omicron Variant. BioRxiv https://doi.org/10.1101/2021.12.16.472934.

Zeng, C., Evans, J.P., Reisinger, S., Woyach, J., Liscynesky, C., Boghdadly, Z. el, Rubinstein, M.P., Chakravarthy, K., Saif, L., Oltz, E.M., et al. (2021b). Impaired neutralizing antibody response to COVID-19 mRNA vaccines in cancer patients. Cell and Bioscience 11. https://doi.org/10.1186/s13578-021-00713-2.

Zeng, C., Evans, J.P., Faraone, J.N., Qu, P., Zheng, Y.M., Saif, L., Oltz, E.M., Lozanski, G., Gumina, R.J., and Liu, S.L. (2021c). Neutralization of SARS-CoV-2 Variants of Concern Harboring Q677H. MBio 12. https://doi.org/10.1128/mBio.02510-21.

Zeng, C., Evans, J.P., King, T., Zheng, Y.M., Oltz, E.M., Whelan, S.P.J., Saif, L.J., Peeples, M.E., and Liu, S.L. (2022). SARS-CoV-2 spreads through cell-to-cell transmission. Proc Natl Acad Sci U S A 119, 1–12. https://doi.org/10.1073/pnas.2111400119.

Zhang, J., Tang, W., Gao, H., Lavine, C.L., Shi, W., Peng, H., Zhu, H., Anand, K., Kosikova, M., Joon Kwon, H., et al. (2022a). Structural and functional characteristics of SARS-CoV-2 Omicron subvariant BA.2 spike. BioRxiv https://doi.org/10.1101/2022.04.28.489772.

Zhang, L., Li, Q., Wu, J., Yu, Y., Zhang, Y., Nie, J., Liang, Z., Cui, Z., Liu, S., Wang, H., et al. (2022b). Analysis of SARS-CoV-2 variants B.1.617: host tropism, proteolytic activation, cell–cell fusion, and neutralization sensitivity. Emerging Microbes & Infections 11, 1024–1036. https://doi.org/10.1080/22221751.2022.2054369.

Zhang, Z., Zheng, Y., Niu, Z., Zhang, B., Wang, C., Yao, X., Peng, H., Franca, D.N., Wang, Y., Zhu, Y., et al. (2021). SARS-CoV-2 spike protein dictates syncytium-mediated lymphocyte elimination. Cell Death and Differentiation 28, 2765–2777. https://doi.org/10.1038/s41418-021-00782-3.

Zhou, D., Dejnirattisai, W., Supasa, P., Liu, C., Mentzer, A.J., Ginn, H.M., Zhao, Y., Duyvesteyn, H.M.E., Tuekprakhon, A., Nutalai, R., et al. (2021). Evidence of escape of SARS-CoV-2 variant B.1.351 from natural and vaccine-induced sera. Cell 184, 2348–2361.e6. https://doi.org/10.1016/j.cell.2021.02.037.

