## Supplementary Figures and legends for "Differential Evasion of Delta and Omicron Immunity and Enhanced Fusogenicity of SARS-CoV-2 Omicron BA.4/5 and BA.2.12.1 Subvariants"

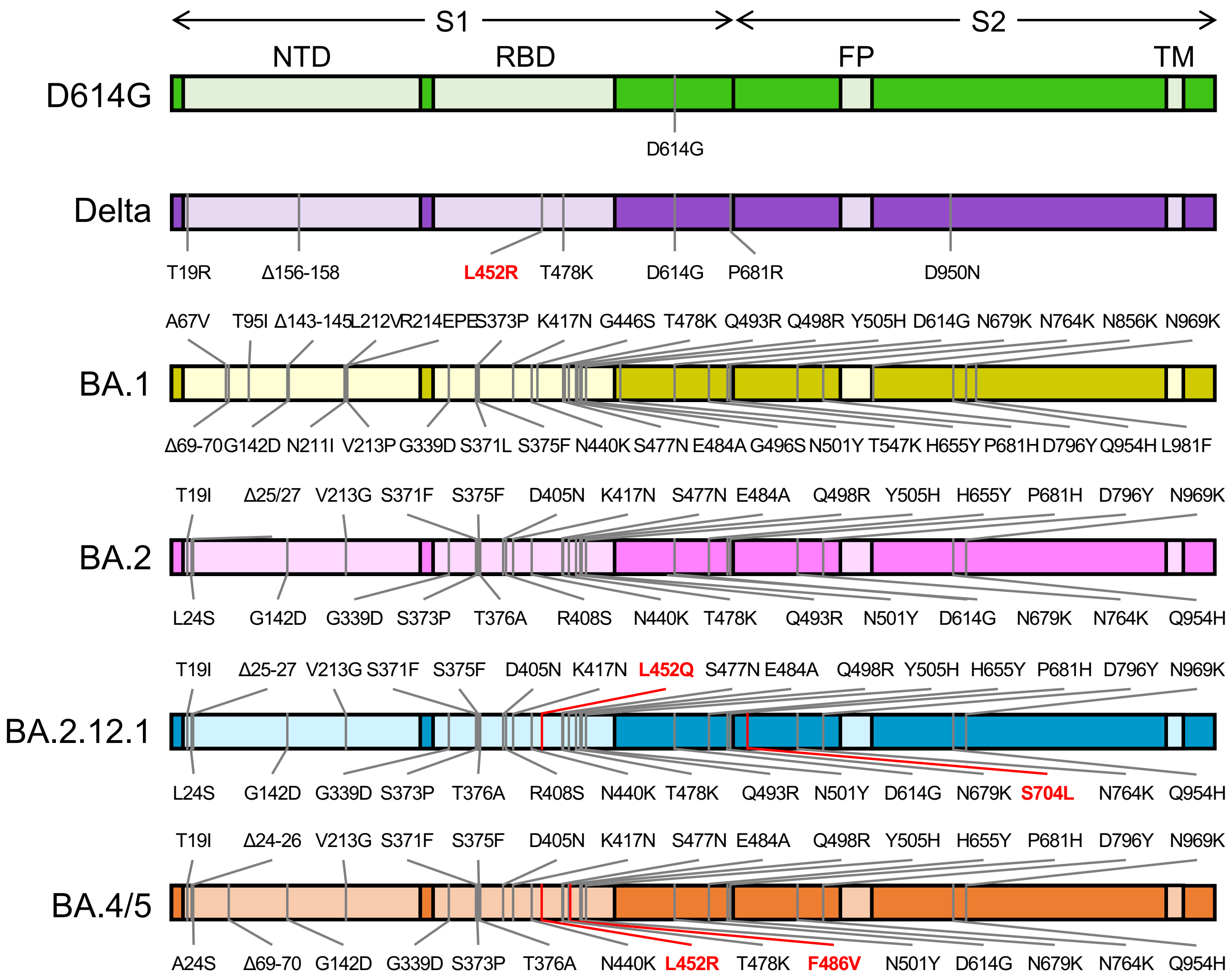


**Figure S1: Schematic of SARS-CoV-2 variant S constructs used for lentivirus pseudotyping.** Diagrams of SARS-CoV-2 S variant constructs used for pseudotyping, which indicate the location of specific mutations as well as the S1 and S2 subunits of S. Domains labeled include the N-terminal domain (NTD), receptor binding domain (RBD), fusion peptide (FP), and transmembrane domain (TM).

Supplementary Figure 1

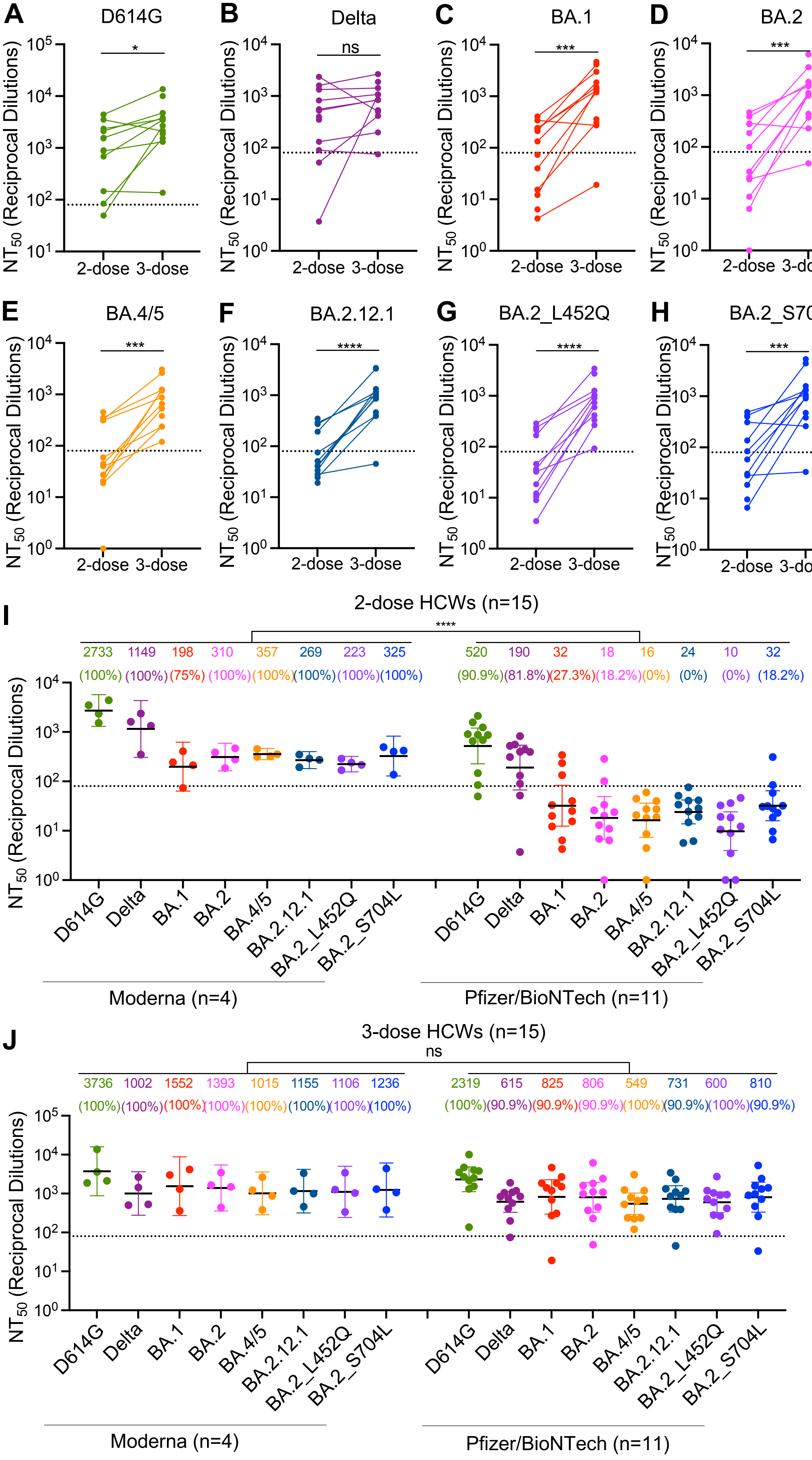


Supplementary Figure 2

**Figure S2: Heterogeneity of neutralization resistance by two-dose and 3-dose HCWs based on individual immunity and vaccine type, corresponds to data in Figure 1.** (**A-H**) Post- second and third dose $NT_{50}$ values plotted pairwise for HCWs for which both time points were analyzed (n=12) against (**A**) D614G, (**B**) Delta, (**C**) BA.1, (**D**) BA.2, (**E**) BA.4/5, (**F**) BA.2.12.1, (**G**) BA.2_L452Q, and (**H**) BA.2_S704L. Significance was determined with a paired two-tailed student's t-test with Welch's correction. (**I**) $NT_{50}$ values for HCWs who received either two doses of Moderna mRNA-1273 (n=4) or Pfizer/BioNTech BNT162b2 (n=11) plotted by vaccine type. (**J**) $NT_{50}$ values for HCWs who received a homologous booster of Moderna (n=4) or Pfizer (n=11) vaccines plotted by vaccine type. $NT_{50}$ values are depicted at the top of graphs (**I and J**) along with the percentage of subjects with $NT_{50}$ values above the limit of detection. Bars represent geometric mean ± 95% confidence interval and significance was determined by two-way repeated measures ANOVA with Bonferroni's multiple testing correction. Throughout p values are represented as $*p < 0.05$, $***p < 0.001$, $****p < 0.0001$, ns, not significant.

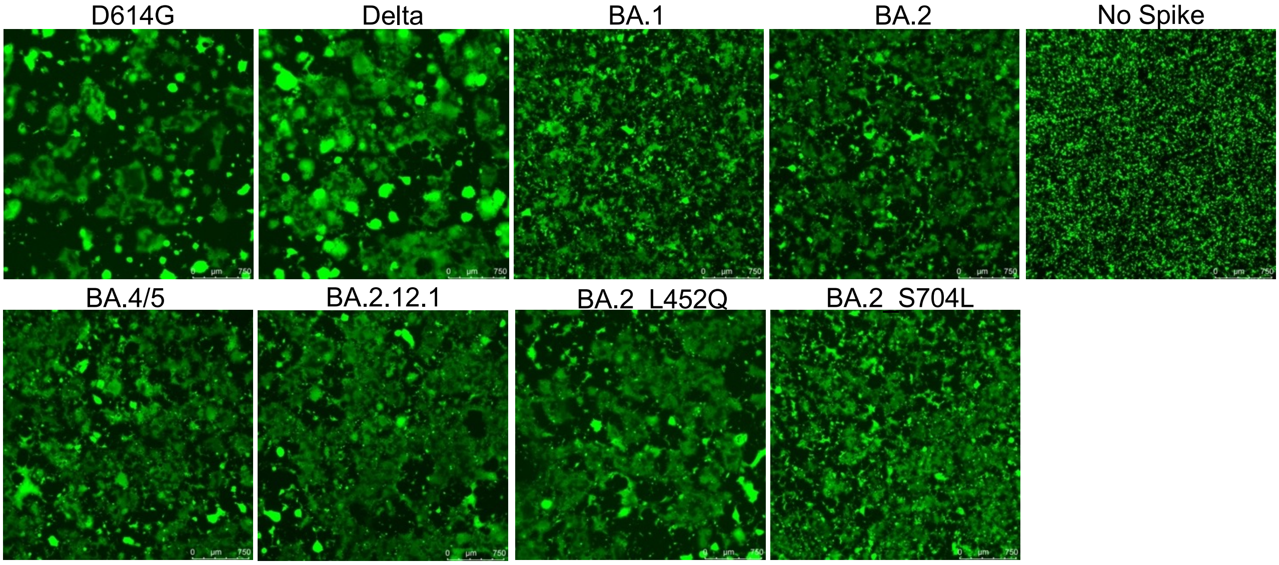


**Figure S3: BA.4/5 and BA.2.12.1 exhibit markedly higher fusion activity than BA.1.** HEK293T cells stably expressing ACE2 were co-transfected with GFP and S constructs and cultured for 48 hrs then imaged to visualize cell-cell fusion.
